# BTBD9 is a novel component of IGF signaling and regulates manganese-induced dopaminergic dysfunction

**DOI:** 10.1101/2021.02.18.431924

**Authors:** Pan Chen, Fuli Zheng, Shaojun Li, Hong Cheng, Julia Bornhorst, Yunhui Li, Bobo Yang, Kun He Lee, Tao Ke, Tanja Schwerdtle, Xiaobo Yang, Aaron B. Bowman, Michael Aschner

## Abstract

Restless legs syndrome (RLS) is a common neurological disorder associated with iron deficiency and dopaminergic (DAergic) neuronal dysfunction. BTBD9 is a genetic risk factor for RLS. However, its molecular function remains largely unknown. Here, we report the interaction between BTBD9, manganese (Mn) and insulin/insulin-like growth factor (IGF) signaling in *Caenorhabditis elegans*, mouse Neuro2a cells and humans. We found that elevated Mn downregulated BTBD9 mRNA levels; in turn, BTBD9 expression attenuated Mn-induced cellular stress and dopaminergic neurodegeneration. As Mn is a known co-factor for insulin receptor and IGF-1 receptor, which activates IGF signaling, we posited that BTBD9 negatively regulates IGF signaling. Our results showed that the protective effects of BTBD9 against Mn toxicity were dependent on the forkhead box O (FOXO) protein. Furthermore, BTBD9 overexpression significantly elevated FOXO level and decreased PKB level, while phosphoinositide-dependent kinase-1 (PDK1) level remained unchanged. We conclude that BTBD9 acts as a key component in the IGF signaling pathway. Meanwhile, the roles of Mn in DAergic neurotoxicity and regulating BTBD9 shed new light on the etiology of RLS.

## Introduction

Restless legs syndrome (RLS) is a common neurological disorder estimated to affect ∼10% of the US population, with symptoms ranging from mild to severe (Allen, Bharmal et al., 2011). RLS is characterized with involuntary twitching leg movements either when asleep or awake (2003), which is associated with sleep deprivation, anxiety, depression and attention-deficit/hyperactivity disorder (ADHD) symptoms (Allen et al., 2011). Moreover, RLS may portend more serious consequences, including hypertension, heart disease and stroke (Li, Walters et al., 2012). The pathobiology of RLS has been linked to deficits in dopaminergic (DAergic) function and iron (Fe) deficiency in the brain (Connor, 2008, Tan, 2006). BTBD9 (BTB domain containing protein 9) has been identified as the most prevalent RLS genetic risk factors for RLS (Stefansson, Rye et al., 2007). It is ubiquitously expressed in both the periphery system and the central nervous system (Lein Hawrylycz et al., 2007). Studies in mouse and fly models have also demonstrated a functional interaction between BTBD9, Fe biology and DAergic activity (DeAndrade, Zhang et al., 2012, Freeman, Pranski et al., 2012).

Manganese (Mn) is an essential nutrient. However, excessive exposure to Mn in occupational and environmental settings may be accompanied by neurotoxicity, and its accumulation in subcortical structures of the basal ganglia that are highly sensitive to oxidative injury, including the substantia nigra, globus pallidus (GP) and the caudate/putamen of the striatum (Olanow, 2004). Chronic childhood exposure to Mn via drinking water preferentially impacts basal ganglia structures (Lao, Dion et al., 2017). The brain area most susceptible to Mn injury is also highly sensitive to oxidative stress, containing metabolically active cell types, particularly tonically active motor neurons that require high levels of ATP for optimal function (Klockgether & Turski, 1989). Recently, a new form of familial parkinsonism was reported in patients carrying homozygous mutations in Mn exporter *SLC30A10*, and all patients exhibited 10-20 fold increase in blood Mn levels concomitant with Mn deposition in the basal ganglia (Quadri, Federico et al., 2012, Tuschl, Clayton et al., 2012). Thus, excess Mn exposure is neurotoxic and likely eliciting DAergic neuronal dysfunction. Mn and Fe have similar chemical and physical characteristics. Mn shares numerous homeostatic and transport pathways with Fe, including the divalent metal transporter (DMT), the transferrin receptor system (TfR) and the Fe exporter, ferroportin (Chen, Chakraborty et al., 2015). It has been demonstrated that Fe deficiency is associated with increased brain deposition of Mn (Au, Benedetto et al., 2008, Garcia, Gellein et al., 2007). These findings indicate a biological basis for RLS and point to a disruption in metal homeostasis, Fe and Mn in particular. A significant concern is whether the cell phenotype, and ultimately the symptoms of RLS patients, are the result of Fe deficiency or elevated concentrations of another metal that opportunistically increases when Fe levels are low. Given the established link between Fe and Mn biology, we deemed it worthwhile to evaluate whether systemic and/or neuronal alterations in Mn may contribute to the etiology of RLS. The insulin/insulin-like growth factor (IGF) signaling pathway has board roles in development and metabolism of the nervous system, including cell proliferation, differentiation, migration, survival, apoptosis, aging and stress response, *etc* (Lewitt & Boyd, 2019). Mn can act as a co-factor for the insulin receptor (IR) and the IGF receptor (IGFR) (Mooney & Green, 1989, Xu, Bird et al., 1995), and the IGF signaling pathway is Mn-regulated (Bryan, Nordham et al., 2020). However,

Here we showed that the RLS risk factor-BTBD9, protected against Mn-induced toxicity, including lethality, oxidative stress, mitochondrial dysfunction and DAergic neurodegeneration. This protection is achieved by regulating the insulin/insulin-like growth factor (IGF) signaling pathway, a pathway known to be regulated by Mn (Bryan et al., 2020, Mooney & Green, 1989, Xu et al., 1995). An etiological role of Mn in RLS would be clinically significant given attempts to treat these patients with Fe supplements (Aurora, Kristo et al., 2012, Ondo, 2010), and if correct, would provide novel translational insights into the biological basis of RLS.

## Results

### Loss of HPO-9 renders worms more susceptible to Mn exposure

*C. elegans* has a single BTBD9 mammalian homolog known as *hpo-9*, with ∼75% sequence similarity (Fig. S1A). A mutant allele *tm3719* (761bp deletion) results in an in-frame deletion, which removes majority of Exon 2. We found this mutant still produced *hpo-9* mRNA (∼1kb), but shorter than the wild type (Fig. S1B). To test our hypothesis, synchronized larva 1 stage nematodes (L1s) were exposed to MnCl_2_. We found that *tm3719* animals were more sensitive to Mn exposure, as indicated by the left shift of the survival curve (Fig. 1A). In contrast, there was no significant difference between the control and *tm3719* animals when exposed to FeCl_2_ or CuCl_2_ (Fig. 1B&C), and the mutant animals showed some tolerance against ZnCl_2_ (Fig. 1D). As the mutants were more sensitive to Mn exposure, we wondered whether this would be reflected in Mn accumulation. However, we did not observe any significant difference between the control and *tm3719* animals after Mn exposure (Fig. 1E), indicating HPO-9 either does not regulate total intracellular Mn content or it is expressed in a limited number of tissues and therefore does not affect overall internal Mn levels. This finding is consistent with our recent report in which RLS patients showed no differences in serum Mn levels by RLS versus control, nor by the BTBD9 genotype (Chen, Bornhorst et al., 2020).

**Figure 1.**
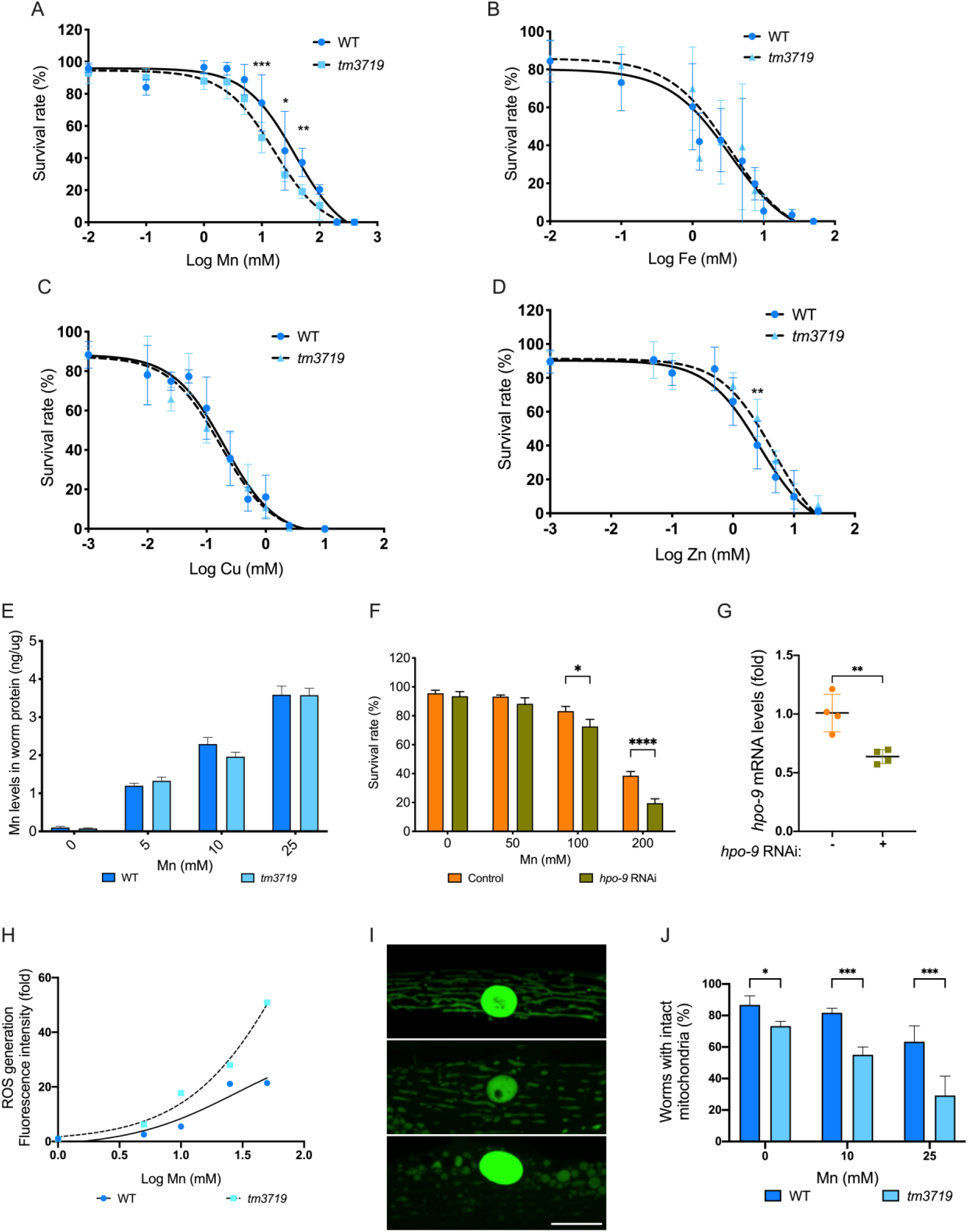
*hpo-9 (tm3719)* mutant were more sensitive to Mn exposure. A-D Survival rate of animals after metal exposure. Synchronized L1 animals were exposed to MnCl2, FeCl2, CuCl2 and ZnCl2, respectively. survival rate was scored two days post exposure. Two-way ANOVA was carried out by Graphpad Prism; *, p<0.05; **, p<0.01; ***, p<0.001; mean±SD (N=6) E Mn levels were determined by ICPMS after exposure. Two-way ANOVA was carried out by Graphpad Prism; mean±SD (N=6). F Survival rates in young adult animals after RNAi feeding and Mn exposure. Two-way ANOVA was carried out by Graphpad Prism; *, p<0.05; ****, p<0.0001; mean±SD (N=9). G hpo-9 mRNA levels in control and knockdown animals were determined by qPCR. Two-way ANOVA was carried out by Graphpad Prism; **, p<0.01; mean±SD (N=4). H ROS levels were determined by measuring DCFDA fluorescence intensity after Mn exposure. Fluorescence intensity was normalized to 0 mM Mn for each strain. Two-way ANOVA shows a significant effect between genotypes (p<0.0001); mean±SD (N=6). I Mitochondrial morphology in Mn-exposed young adult nematodes. Top, intact; middle, intermediate; bottom, fragmented. Scale bar, 10 μM. J Quantitative analysis of mitochondrial morphology by scoring worms with intact mitochondria. Two-way ANOVA was carried out by Graphpad Prism;*, p<0.05, ***, p<0.001; mean±SD (N=3).

To confirm the results from the mutant study, we knocked down *hpo-9* using RNAi feeding and exposed these animals to Mn. A RNAi sensitive strain GR1373 were used. L1 animals were fed with bacteria carrying either L4440 empty vector (control) or dsRNA for *hpo-9* (RNAi) until they reached young adult stage. After Mn exposure, we found that knockdown of *hpo-9* significantly decreased survival rate at 100 and 200 mM of Mn exposure, compared with the control (Fig. 1F). To confirm that the RNAi was specifically targeting *hpo-9*, qPCR was performed to quantify *hpo-9* mRNA levels in control and knockdown animals. *hpo-9* level was knocked down by ∼40% in nematodes fed with RNAi bacteria, compared with the L4440 control (Fig. 1G), ensuring the efficacy of the RNAi.

### Elevated reactive oxygen species (ROS) and mitochondrial dysfunction in *hpo-9* mutant upon Mn exposure

Mn exposure promotes generation of ROS, which results in elevated oxidative stress and mitochondrial dysfunction (Fernsebner, Zorn et al., 2014, Malecki, 2001). Here we used a fluorescent probe 2′,7′-dichlorofluorescein diacetate (DCFDA) to determine ROS levels in the nematodes exposed to Mn. The non-fluorescent DCFDA can be rapidly oxidized to a highly fluorescent 2′,7′-dichlorofluorescein (DCF) in the presence of ROS (Yoon, Lee et al., 2018). We found that Mn exposure resulted in elevated fluorescence intensity in a dose-dependent pattern in both control and the mutant animals (Fig. 1H), indicating Mn exposure promotes generation of ROS. In addition, *hpo-9* mutants showed consistently higher fluorescence intensity when compared with the control (Fig. 1H). These results suggest that Mn induces higher oxidative stress in *hpo-9* mutants than WT worms and that HPO-9 protects nematodes against Mn-induced toxicity by downregulating Mn-dependent ROS production.

RLS is associated with impairment of mitochondrial function and reduced respiratory capacity (Haschka, Volani et al., 2019). As Mn is known to accumulate in the mitochondria (Gavin, Gunter et al., 1992) and cause oxidative stress and mitochondrial dysfunction (Fernsebner et al., 2014, Malecki, 2001), we tested whether HPO-9 protected mitochondria from Mn exposure. The strain SD1347 (*myo-3p*::GFP::LaZ::NLS + *myo-3p*::mitoGFP) with GFP labelled mitochondria and nucleus in the body wall muscle cells, was crossed with the *tm3719* mutant to generate the MAB506 strain. Young adult stage animals (SD1347 and MAB506) were exposed to Mn for 2 hours and allowed to recover for another 2 hours before imaging. We then examined mitochondrial morphology in the nematode. The morphological categories were defined as previously described (Regmi, Rolland et al., 2014) with slight modifications: 1) images with a majority of long interconnected tubular mitochondrial networks were defined as “intact” (Fig. 1I); 2) images with a combination of interconnected and some smaller fragmented mitochondria networks were classified as “intermediate” (Fig. 1J); 3) images with a majority of short, sparse round or a collapse of muscular mitochondria were classified as “fragmented” (Fig. 1K). The intermediate and fragmented mitochondria were considered as defective. We scored the percentage of worms with intact mitochondria network and found that the mutant animals unexpectedly showed a slight but significant decrease (∼15%, p=0.0212) in intact mitochondria even without Mn exposure, compared with the WT control (Fig. 1L), indicating BTBD9 regulates mitochondrial homeostasis. After Mn exposure, the *tm3719* mutant consistently showed lower ratio of intact mitochondria than the WT. These results suggest that HPO-9 likely plays an important role, not only on maintaining mitochondrial morphology at normal physiological conditions, but also in protecting against Mn-induced oxidative stress.

### Mn exposure down-regulates *hpo-9* mRNA levels

As knocking down of *hpo-9* rendered animals more sensitive to Mn, next, we determined whether *hpo-9* levels were lower in the mutant carrying *tm3719* allele. Nematodes at L1 and young adult stages were analyzed in the absence of Mn exposure. To detect the gene product of *tm3719* allele (with deletion in Exon 2), a Taqman probe with Exon 4-5 boundary was selected for q-PCR. We found a significant decrease (∼50%) of *hpo-9* mRNA levels in the mutant at both L1 and young adult stages (Fig. 2A), indicating the *tm3719* allele is deficient in gene transcription. Interestingly, Mn exposure resulted in a significant decrease (∼40%) in *hpo-9* mRNA in the wild type animals, but not the mutants (Fig. 2B). In contrast, The mRNA levels remained unchanged in in the mutant across different doses of Mn exposure.

**Figure 2.**
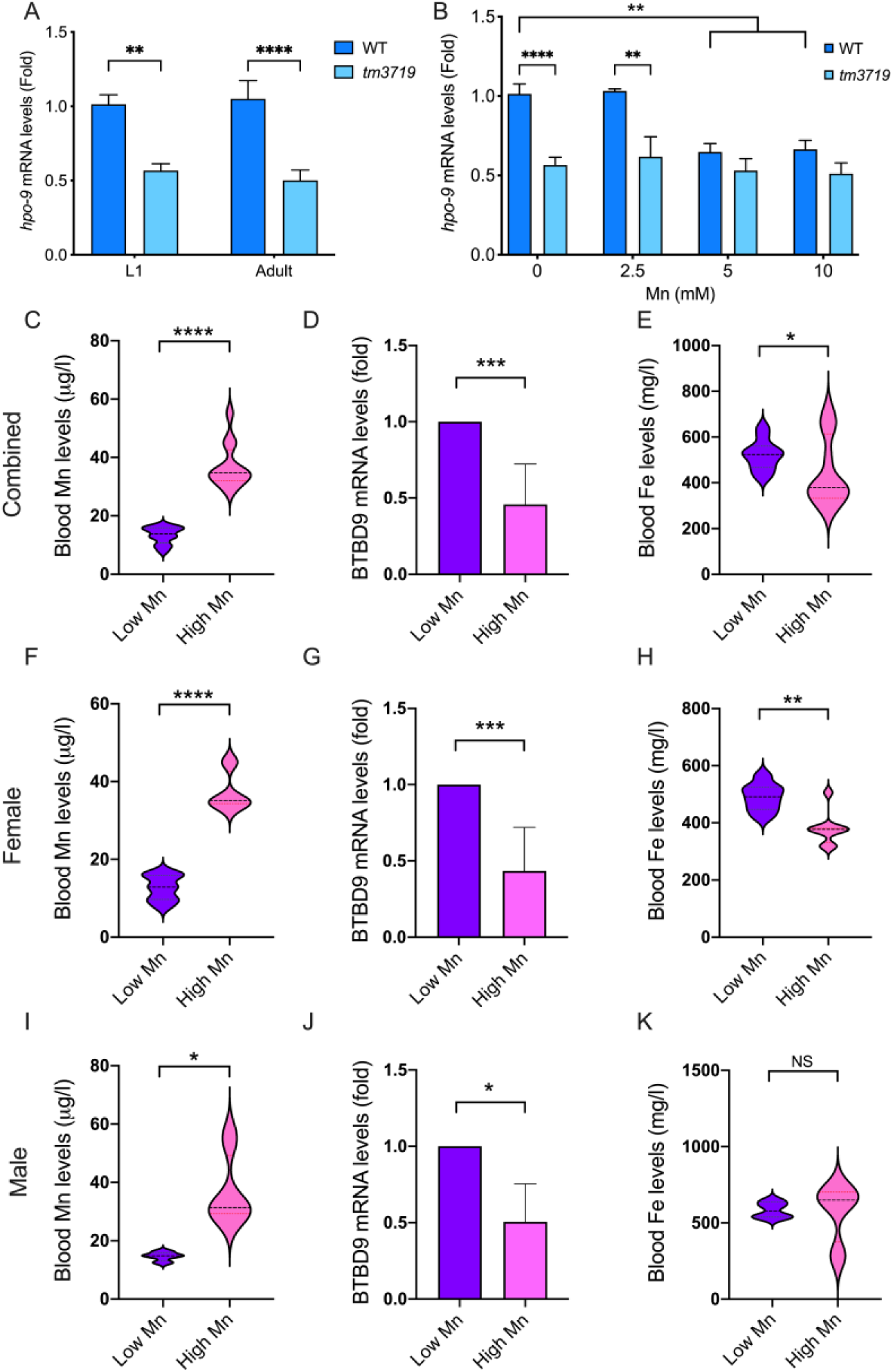
Mn exposure downregulated *hpo-9*/BTBD9 mRNA levels in both *C. elegans* and human blood samples. A *hpo-9* mRNA levels in L1 larva and young adult nematodes determined by RT-PCR without Mn exposure. Two-way ANOVA was carried out by Graphpad Prism;**, p<0.01, ****, p<0.0001; mean±SD (N=8). B *hpo-9* mRNA levels in L1 larva of WT and *tm3719* animals after Mn exposure. All *hpo-9* levels were normalized to the control animals at 0 mM Mn. Two-way ANOVA was carried out by Graphpad Prism;**, p<0.01, ****, p<0.0001; mean±SD (N=8). C,F&I Blood Mn levels in in combined (C), female (F) and male (I) blood samples quantified by ICPMS. *t* test was carried out by Graphpad Prism; *, P<0.05, ****, p<0.0001; mean±SD (C, n=12; F, n=8, I, n=4). D,G&J *BTBD9* mRNA levels in combined (D), female (G) and male (J) blood samples quantified by qPCR. *t* test was carried out by Graphpad Prism; *, P<0.05, ***, p<0.001; mean±SD (D, n=12; G, n=8, J, n=4). E,H&K Blood Fe levels in in combined (C), female (F) and male (I) blood samples quantified by ICPMS. *t* test was carried out by Graphpad Prism; *, P<0.05, **, p<0.01, NS, non-significant; mean±SD (C, n=12; F, n=8, I, n=4).

To corroborate this observation in humans, blood samples from 24 individuals were collected and analyzed. These age- and sex-matched individuals were separated into two groups (12 each, including 8 women and 4 men) defined as Control-Low Mn (with Mn levels ranging from 7.9-16.5 μg/l) and High Mn (with Mn levels ranging from 28.8-55.2 μg/l) (Fig. 2C, Table 1). We found that individuals with higher whole blood Mn levels had a significantly lower BTBD9 mRNA levels (∼50%) (Fig. 2D), consistent with the results in the nematode. Given that the major risk genotypes (e.g. single nucleotide polymorphisms (SNPs)) of BTBD9 associated with RLS risk are found in the noncoding regions of the gene (Jiménez-Jiménez, Alonso-Navarro et al., 2018, Vilariño-Güell, Chai et al., 2009, Winkelmann, Schormair et al., 2007) and thus are not thought to alter the protein sequence, it is plausible that those SNPs regulate the transcription of BTBD9 and alter its mRNA levels. Interestingly, in people with higher blood Mn, Fe levels were significantly decreased (Fig. 2E). In the individual with the highest Mn level (55.2 μg/l), whole blood Fe levels were only about half of the control group (284 vs. 524 mg/l) and the BTBD9 mRNA level was about 16% of control (Table 1). As the prevalence of RLS in women is much higher (almost double) than in men (Tison, Crochard et al., 2005), we compared the sex-dependent effect on BTBD9 and Fe levels. In both men and women, high whole blood Mn led to lower BTBD9 levels (Fig. F,G,I&J). Notably, high Mn was associated with significantly lower Fe levels only in women, but not in men (Fig. 2H&K), suggesting a sex-dependent pattern of Fe levels may exist. In addition, the results indicate that BTBD9 transcription is specifically associated with Mn levels, rather than Fe levels. Together, these results indicated that excessive Mn is able to down regulate *BTBD9*/*hpo-9* expression, at least at transcriptional level; as a result, decreased *BTBD9*/*hpo-9* possibly results in susceptibility to Mn-induced toxicity.

**Table 1.** Mn and Fe levels in the blood of Age and Sex matched groups. (F, female; M, male; Work Years, years working in the factory).

| Group | Sex | Age | Work Years | Mn (µg/l) | Fe (mg/l) | Relative BTBD9 mRNA level |
| --- | --- | --- | --- | --- | --- | --- |
| 1 | F | 49 | 31 | 10.1 | 462.1 | 1.00 |
|  | F | 52 | 33 | 32.9 | 506.2 | 0.70 |
| 2 | F | 50 | 30 | 13.0 | 525.5 | 1.00 |
|  | F | 50 | 34 | 35.4 | 372.2 | 0.26 |
| 3 | F | 49 | 24 | 9.5 | 418.7 | 1.00 |
|  | F | 48 | 29 | 45.0 | 377.2 | 0.17 |
| 4 | F | 41 | 24 | 15.9 | 499.7 | 1.00 |
|  | F | 44 | 25 | 34.4 | 381.2 | 0.90 |
| 5 | F | 45 | 25 | 15.9 | 565.9 | 1.00 |
|  | F | 44 | 25 | 34.4 | 381.2 | 0.12 |
| 6 | F | 50 | 24 | 16.5 | 522.0 | 1.00 |
|  | F | 46 | 26 | 35.2 | 318.5 | 0.35 |
| 7 | F | 48 | 29 | 12.8 | 483.0 | 1.00 |
|  | F | 48 | 29 | 45.1 | 377.1 | 0.67 |
| 8 | F | 48 | 23 | 7.9 | 442.6 | 1.00 |
|  | F | 46 | 26 | 35.2 | 318.5 | 0.30 |
| 9 | M | 52 | 34 | 12.5 | 610.2 | 1.00 |
|  | M | 55 | 36 | 30.8 | 719.1 | 0.69 |
| 10 | M | 55 | 31 | 14.9 | 544.3 | 1.00 |
|  | M | 57 | 36 | 28.8 | 648.9 | 0.49 |
| 11 | M | 47 | 23 | 14.7 | 645.7 | 1.00 |
|  | M | 48 | 22 | 31.8 | 651.6 | 0.68 |
| 12 | M | 56 | 36 | 16.3 | 535.5 | 1.00 |
|  | M | 56 | 36 | 55.2 | 283.7 | 0.16 |

### Expression pattern of *hpo-9* in *C. elegans*

Given that BTBD9 is expressed in the DAergic neurons and RLS patients are associated with DAergic neurological dysfunction (Freeman et al., 2012), next, we determined whether *hpo-9* is expressed in DAergic neurons of the nematodes and functions to maintain normal DAergic activities. A transcriptional GFP reporter strain MAB336 (*hpo-9p::*GFP) was generated by using *hpo-9* promoter sequence (∼1.5kb upstream of the start codon) to drive GFP expression. We consistently observed a ubiquitous expression from larva 1 (L1) to adult stage, despite mosaicism due to the extra-chromosomal array, indicating HPO-9 was expressed throughout development in *C. elegans.* More specifically, HPO-9 expression was observed primarily in the head neurons and pharynx, and also but weaker in the posterior intestine, body wall muscle cells and hypodermal seam cells (Fig. 3A&B).

**Figure 3.**
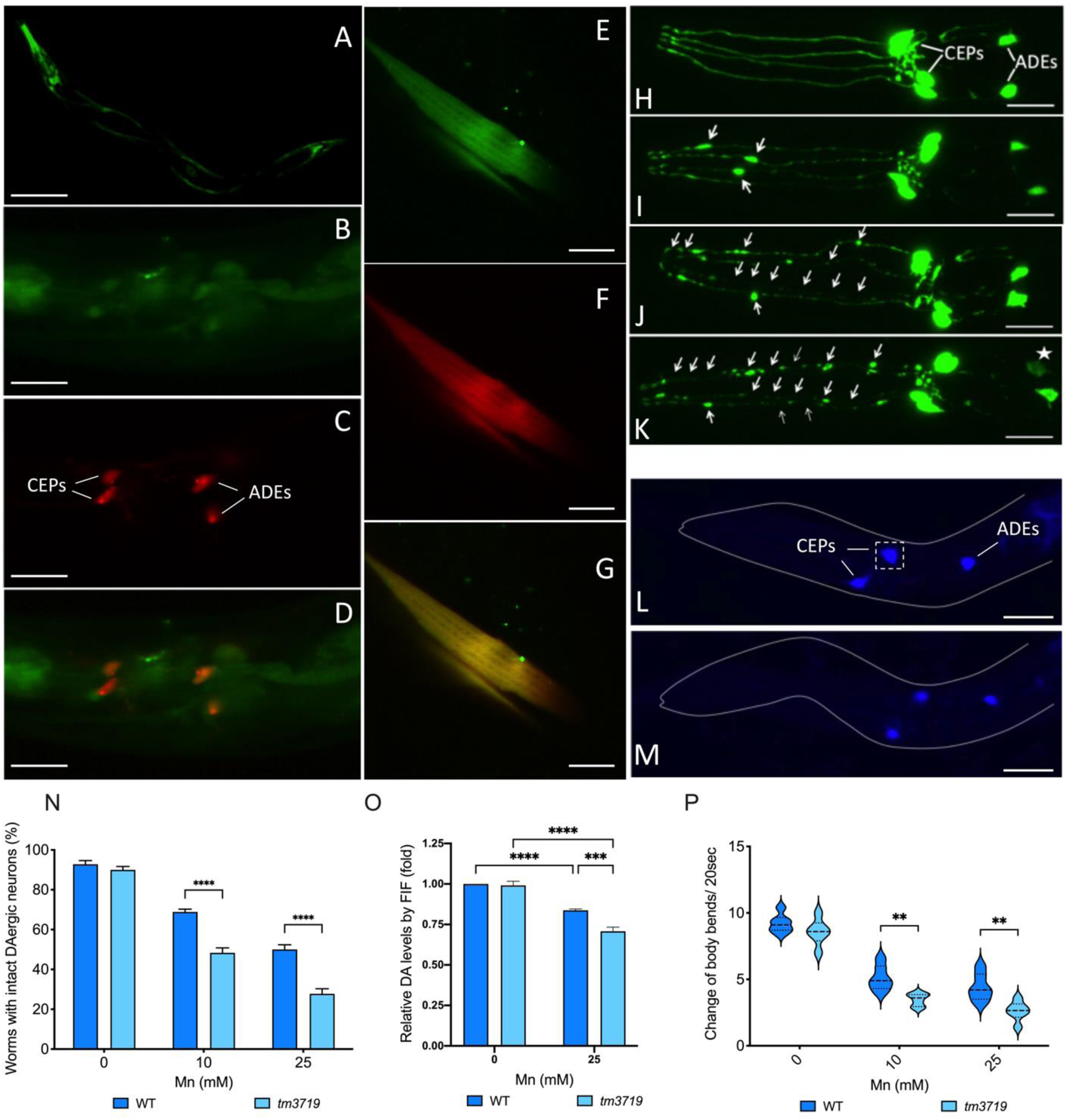
HPO-9 expression pattern and Mn-induced neurotoxicity in the nematode. A Expression pattern of the transcriptional GFP resporter *hpo-9p::*GFP in a Larva 3 (L3) stage animal. Scale bar, 100 μM. B-D *hpo-9* in DAergic neurons. B, *hpo-9p*::GFP expression in the head; C, mCherry labeled DAergic neurons; D, merged GFP and mCherry. Scale bars, 15 μM. E-G Cytosolic localization of HPO-9 protein. E, intracellular expression of a translational GFP reporter (*unc-*54::HPO-9::GFP) in the body wall muscle cells; F, DsRed protein expressed in the cytoplasm; G, merged GFP and DsRed. Scale bars, 15 μM. H-K DAergic neurons in MAB300 worms exposed to 0 mM (H) or 10 mM MnCl_2_ for 1 h (I– K). H, intact DAergic neurons in the head, including 4 CEPs and 2 ADEs. I–K, impaired DAergic neurons, arrowheads indicate damaged neuronal processes, asterisk indicates alternations in cell bodies. Scale bars, 10 μM. L&M DA staining by Formaldehyde induced fluorescence (FIF) in worms exposed to 0 mM (L) and 25mM (M) MnCl_2_. The dash square indicate the region with individual DAergic neurons for measuring fluorescence intensity. Scale bars, 10 μM. N Quantitative analysis of DAergic neuron morphology after Mn exposure. Worms were recovered for 2 hours before analysis. Two-way ANOVA was carried out by Graphpad Prism; ****, p<0.0001; mean±SD (N=9). O Quantitative analysis of dopamine levels by FIF. After Mn exposure, animals were recovered for 2 hours before paraformaldehyde staining. The fluorescence intensity was normalized to WT animals at 0 mM Mn. Two-way ANOVA was carried out by Graphpad Prism;***, p<0.001, ****, p<0.0001; mean±SD (N=3). P Quantitative analysis of basal slowing response by scoring body bends in 20 seconds. After Mn exposure, worms were grown to L4 stage for the behavioral assay. Two-way ANOVA was carried out by Graphpad Prism;**, p<0.01; mean±SD (N=6).

*C. elegans* hermaphrodites have 8 DAergic neurons, including 4 CEPs and 2 ADEs in the head (Fig. 3C&H) and 2 PDEs in the posterior, which can be visualized by *dat-*1 (DA transporter) promoter driven mCherry or GFP. To study whether *hpo-9* is expressed in DAergic neurons, we crossed the transcriptional reporter strain MAB336 with OH7193 (*dat-1p::*mCherry, *ttx-3p::*mCherry), which has mCherry labeled DAergic neurons (Fig. 3C). Using confocal microscopy, we found that the GFP signal was co-localized with *dat-1p*::mCherry labeled DAergic neurons (MAB337, Fig. 4B-D), indicating *hpo-9* was expressed in DAergic neurons. Moreover, a translational reporter strain MAB335 (*unc-54p::*HPO-9::GFP, *myo-3*::DsRed) was created to determine its intracellular localization in the body wall muscle cells. We found that HPO-9::GFP was confocalized with DsRed, a red fluorescent protein constantly expressed in the cytoplasm (Fig. 4E-G). The results indicated that HPO-9 was localized in the cytosol, consistent with the previously findings in the human and *Drosophila* (Freeman et al., 2012). Taken together, these results suggest that HPO-9 is a cytoplasmic protein primarily expressed in the pharynx and head neurons, however, its expression is weak in other soma tissues, which may explain why no difference in internal metal levels was observed in the mutant animals (Fig. 1E). In addition, HPO-9 is expressed in the DAergic neurons, suggesting it plays a role in these neurons.

**Figure 4.**
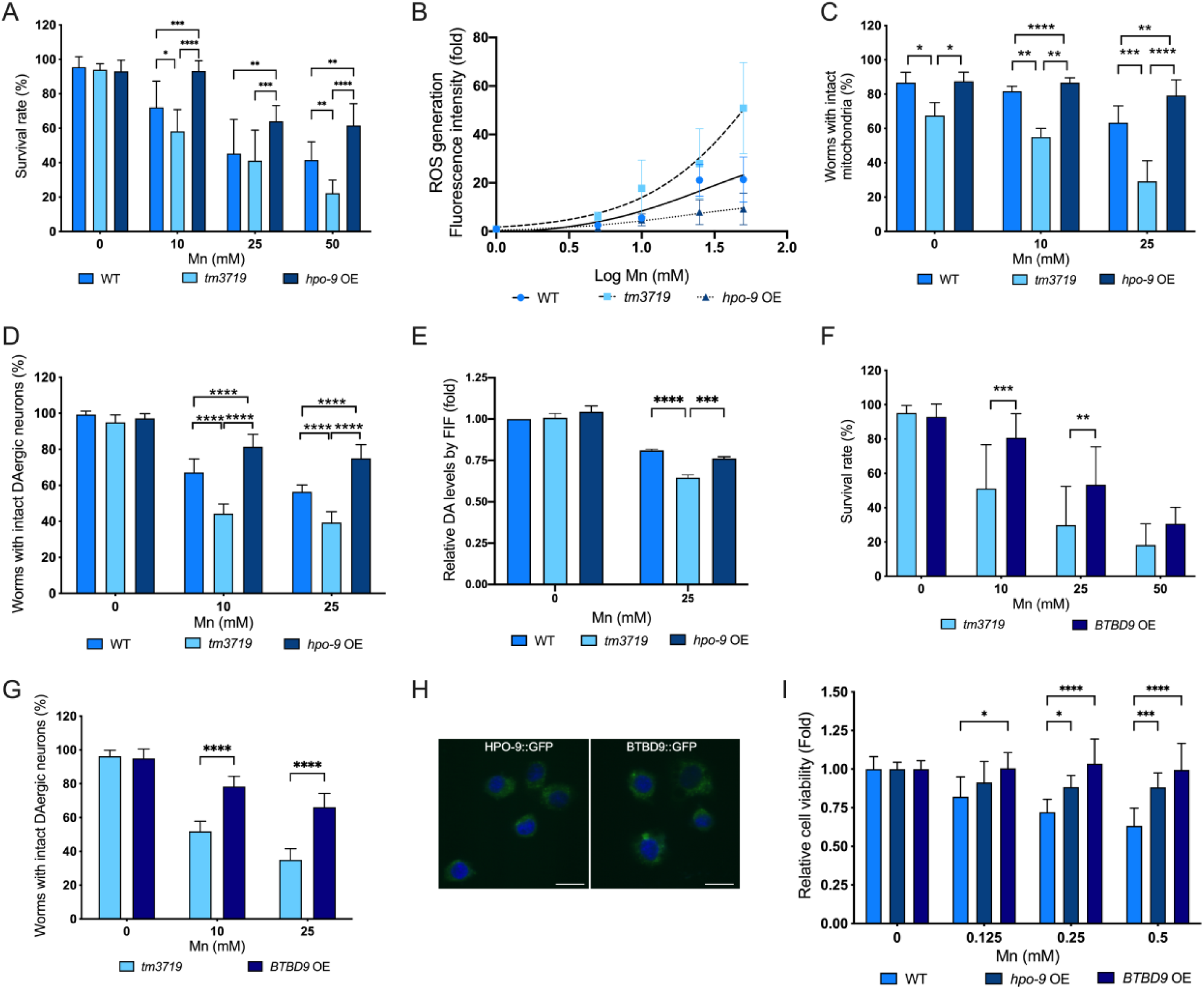
Expression of *hpo-9* in the *tm3719* animals restores the mutant defects. A&F Survival rate of L1s after Mn exposure. Two-way ANOVA was carried out by Graphpad Prism; *, p<0.05, **, p<0.01,***, p<0.001, ****, p<0.0001; mean±SD (N=9). B ROS levels measured by DCFDA assay. Fluorescence intensity was normalized to 0 mM Mn for each strain. Two-way ANOVA by Graphpad Prism shows a significant effect between genotypes (p<0.0001); mean±SD (N=6). C Quantitative analysis of mitochondrial morphology. Two-way ANOVA was carried out by Graphpad Prism; *, p<0.05, **, p<0.01,***, p<0.001, ****, p<0.0001; mean±SD (N=6). D&G Quantitative analysis of DAergic neuron morphology after Mn exposure. Two-way ANOVA was carried out by Graphpad Prism; ****, p<0.0001; mean±SD (N=7). E Dopamine levels measured using FIF. The fluorescence intensity was normalized to WT animals at 0 mM Mn. Two-way ANOVA was carried out by Graphpad Prism; ***, p<0.001, ****, p<0.0001; mean±SD (N=3). H Intracellular localization of GFP tagged HPO-9 and BTBD9 in Neuro2A cells. Green, GFP tagged proteins in the cytosol; blue, nucleus by DAPI staining. Scale bars=15 µM. I Cell viability by MTT assay. Two-way ANOVA was carried out by Graphpad Prism; *, p<0.05,***, p<0.001, ****, p<0.0001; mean±SD (N=6).

### Loss of HPO-9 resulted in susceptibility to Mn-induced neurotoxicity

As HPO-9 is present in DAergic neurons, we expected that it would protect DAergic neurons from Mn-induced neurotoxicity. Exposure of worms to Mn induced morphological changes characteristic of neurodegeneration in the 6 head DAergic neurons, causing puncta, blebs, neuronal shrinkage and loss of cell bodies (Fig. 3I-K) (Leyva-Illades, Chen et al., 2014). These morphological changes represented phenotypes of neurodegeneration, which were confirmed by transmission electronic microscopy (TEM) (Nass, Hall et al., 2002). Nematodes expressing *dat-1p*::GFP (MAB300) were crossed with *hpo-9* mutant to generate MAB382 strain. Animals were synchronized at L1 stage, followed by Mn exposure. Under normal condition in the absence of excess Mn exposure, we found no difference between the control and mutant worms in the morphology of DAergic neurons. However, *tm3719* worms had a significantly lower percentage of morphologically normal DAergic neurons than control worms upon Mn exposure (Fig.3N), indicating loss of HPO-9 leads to increased susceptibility to Mn-induced neurotoxicity. We also measured dopamine (DA) levels in these animals using formaldehyde induced fluorescence (FIF) technique. In *C. elegans*, DAergic neurons can be stained by paraformaldehyde solution and the generated fluorescence intensity represents DA levels (Fig. 3L&M) (Lints & Emmons, 1999, Rivard, Srinivasan et al., 2010). After Mn exposure and recovery, L1 animals were stained by FIF and the fluorescent images were taken by confocal microscopy. We found that Mn exposure decreased DA levels in both WT and tm3719 animals, while tm3719 worms showed significantly lower DA levels than WT worms (Fig. 3O). To determine if these changes were associated with DAergic dysfunction, we performed a behavioral assay to examine the functional outcome of Mn exposure. Well-fed N2 (WT control) worms slow down their locomotion when encountering a bacterial lawn (basal slowing response), a behavior that is exclusively controlled by DAergic neurons in *C. elegans* (Sawin, Ranganathan et al., 2000). This behavior can be quantified by counting the changes in the number of body bends over a defined time on plates with or without bacteria (Sawin et al., 2000). *tm3719* worms showed no difference in this behavior without Mn exposure, while Mn exposure caused a significant decrease in basal slowing response in the mutant animals, when compared with the control (Fig. 3P). Taken together, these results showed that: 1) under normal physiological conditions, loss of HPO-9 did not result in DAergic neuronal defect; in the presence of Mn, *tm3719* worms had a significant difference in DAergic morphology changes, DA levels and behavior defects compared with the control, indicating the mutants were more sensitive to Mn induced neurotoxicity. Thus, HPO-9 protects against Mn-induced neurotoxicity. As Mn level was not altered (Fig. 1E), the protection is possibly achieved through modulating oxidative stress induced by Mn exposure.

### Expression of *hpo-9* in the *tm3719* animals restores the mutant defects

To further confirm the protection afforded by HPO-9 against Mn-induced DAergic neurodegeneration, and to exclude the possibility that defects in *tm3719* were caused by other mutations, we expressed the WT *hpo-9* gene in the *tm3719* animals to study whether the defects could be rescued. The stain MAB415 (*eft-3p*::*hpo-9::FLAG*; *hpo-9(tm3719)*) was generated by microinjection with HPO-9 expressed in all somatic tissues. Expression of HPO-9 was confirmed by western blot analysis (Fig. S2A). We found that transgenic over-expression of *hpo-9* (OE) itself (0 mM) had no effect on worm survival; in the presence of Mn, it not only recovered the lethality defect in the mutant, but also empowered more resistance to Mn than in the control (Fig 4A). We also determined the ROS levels in these strains using DCFDA as previously described (Fig.1H). In the presence of Mn, the ROS levels in the *hpo-9* OE worms were significantly lower than both the control and *tm3719* animals (Fig. 4B). These data indicate that expression of *hpo-9* decreased ROS production caused by Mn exposure. Next, we wondered whether *hpo-9* OE restores mitochondrial morphology defect in *tm3719* animals. MAB509 strain was generated by crossing MAB415 with SD1347 strains. We examined the mitochondrial morphology in SD1347, MAB506 and MAB509 animals as described in Fig. 1I-L. In the absence of Mn, *hpo-9* OE animals had normal mitochondria, indicating expression of *hpo-9* rescued the morphological defect in the mutant (Fig. 4C). After Mn exposure, *hpo-9* OE showed significantly higher percentage of intact mitochondria than both the control and mutant.

To validate HPO-9 function in the DAergic neurons, a strain MAB414 (*dat-1p*::*hpo-9::FLAG*; *hpo-9(tm3719)*) was generated with HPO-9 exclusively expressed in DAergic neurons, rather than the whole animal. Expression of *hpo-9* was confirmed by semi-quantitative RT-PCR (Fig. S2C), due to the limited expression in only 8 neurons. We found that overexpression of *hpo-9* (OE) did not alter DAergic morphology (0 mM); upon Mn exposure, *hpo-9* (OE) animals had significantly higher percentage of WT DAergic neurons than both mutant and control worms (Fig. 4D). Dopamine levels were also determined by FIF. We found that *hpo-9* (OE) animals had significantly elevated DA levels than *tm3719* mutants, similar to the WT animals (Fig. 4E). As HPO-9 was only present in DAergic neurons, the protection was directly from HPO-9 in the DAergic neurons, rather than other somatic tissues, excluding the possibility that the protection was from the overall effect of HPO-9 expression in the whole animal (Fig. 4A). Taken together, expression of *hpo-9* in the mutant fully recovered the defects, confirming its protective function against Mn-induced oxidative stress and mitochondrial dysfunction.

Next, we determined whether the human homolog BTBD9 shared the same function as HPO-9. BTBD9 has several isoforms, here we selected isoform a in the following study. Similarly, we created MAB425 (*eft-3p*::*BTBD9::FLAG*; *hpo-9(tm3719)*) and MAB405 (*dat-1p*::*BTBD9::FLAG*; *hpo-9(tm3719)*) with BTBD9 expressed in the whole animal and DAergic neurons (Fig. S2B&D), respectively. We found that expression of BTBD9 significantly increased survival rate in the mutant background upon Mn exposure (Fig. 4F); BTBD9 OE also protected DAergic neurons from Mn-induced morphological changes (Fig. 4G). Taken together, our results suggested that HPO-9 and BTBD9 share the same function in protecting against Mn-induced lethality and neurotoxicity.

### BTBD9/*hpo-9* protects against Mn-induced toxicity in Neuro2A cells

Given the protective role of BTBD9/*hpo-9* in *C. elegans*, we assumed that it has the similar function in mammalian systems. We expressed GFP tagged *hpo-9* and BTBD9 isoforms a in mouse neuroblastoma Neuro2A cells. Both HPO-9 and BTBD9a were present in the cytoplasm (Fig. 4H), consistent with the findings in *C. elegans* (Fig. 3). Mn exposure decreased the cell survival rate in control cells; in contrast, the presence of HPO-9 and BTBD9 significantly increased cell viability (Fig. 4I), consistent with the results in *C. elegans*.

### HPO-9 regulates Fe levels rather than Mn

As *hpo-9* re-expression attenuated Mn-induced toxicity, we first assumed that it might downregulate Mn levels in the nematode. Previously, we failed to find differences between the control and *tm3719* animals, which might be due to the limited expression in worms (Fig. 3A). Now, with whole tissue expression of *hpo-9* in MAB415 worms, we expected to see a significant decrease of internal Mn levels. However, this was not the case as the *hpo-9* OE strain had similar internal Mn levels as the control and *tm3719* animals (Fig. 5A). This result suggests that HPO-9 alleviates Mn-induced toxicity not by regulating Mn transport. Interestingly, we found that Fe levels were significantly elevated in *hpo-9* OE worms, compared with both the control and *tm3719* animals, regardless of Mn exposure (Fig. 5B). This result is consistent with the fact that Fe levels are lower in the serum and brains of RLS patients, and corroborates the clinical study of Fe supplementation in reducing RLS symptoms (Aurora et al., 2012, Ondo, 2010).

**Figure 5.**
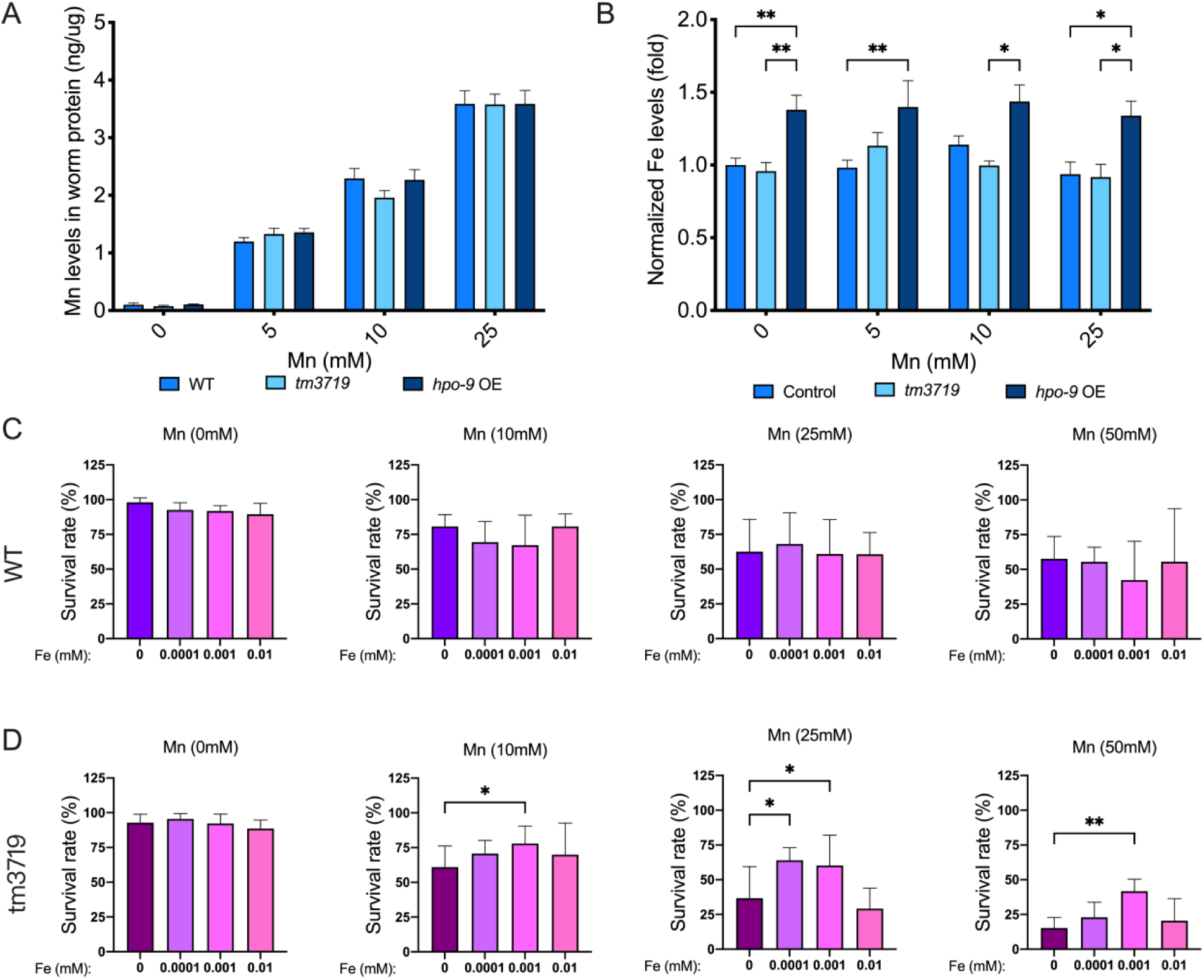
Mn and Fe levels in treated animals and the protective effect of Fe supplementation in in *tm3719* worms. A Mn concentrations measured by ICP-MS. Two-way ANOVA was carried out by Graphpad Prism; no significant difference was observed; mean±SD (N=6). B Fe concentrations measured by ICP-MS. Two-way ANOVA was carried out by Graphpad Prism; *, p<0.05,**, p<0.01; mean±SD (N=6). C Survival of WT animals with Fe supplementation before Mn exposure. Synchronized L1 animals were pre-incubated with FeCl_2_ (0-0.01mM) for 30 min. Fe was then washed and worms were exposed to Mn as described in Fig. 1A. One-way ANOVA was carried out by Graphpad Prism; no significant difference was observed; mean±SD (N=9). D Survival of tm3719 animals with Fe supplementation before Mn exposure. One-way ANOVA was carried out by Graphpad Prism; *, p<0.05; **, p<0.01; mean±SD (N=9).

### Non-toxic Fe treatment protected *tm3719* animals against Mn-induced toxicity

Fe deficiency is commonly observed in the brain of RLS patients (Connor, 2008) and Fe supplementation is clinically effective in reducing RLS symptoms (Aurora et al., 2012, Ondo, 2010). Combined with our findings (Fig. 5B), we wondered whether pre-incubation of *tm3719* animals with low levels of Fe would protect worm upon Mn exposure. To test this hypothesis, L1 animals were pre-incubated with FeCl_2_, followed by Mn exposure. We found that exposure low levels of FeCl_2_ (0.0001 and 0.001 mM) significantly increased *C. elegans* survival rate upon later Mn exposure (Fig. 5G,H,I&J) in *tm3719* animals. In contrast, Fe pre-incubation had no effect in wild type control animals (Fig. 5C,D,E&F). This result recapitulated the effect of Fe supplementation in clinical studies in RLS patients (Aurora et al., 2012, Ondo, 2010).

### Protective effects of HPO-9 are dependent on FOXO/DAF-16

As HPO-9 does not regulate internal Mn concentrations, it is likely that HPO-9 functions to attenuate Mn-induced oxidative stress and mitochondrial dysfunction. It is known that Mn can act as a co-factor for the insulin receptor (IR) and the IGF receptor (IGFR) (Mooney & Green, 1989, Xu et al., 1995), and that the IGF signaling pathway is Mn-regulated (Bryan et al., 2020). Previously, we have shown that loss-of-function mutants of AKT serine/threonine kinase (*akt-1* and *akt-2*) are more resistant to Mn induced lethality, although they have similar internal Mn levels, when compared with WT N2 worms (Peres, Arantes et al., 2018). Meanwhile, increased FOXO level and mutation in IGFR recovered lifespan reduction and dauer movement defects caused by Mn exposure, respectively (Chen, DeWitt et al., 2015). Therefore, we tested whether HPO-9 acts via DAF-16/FOXO signaling. To test this hypothesis, we first analyzed survival rate of the control (WT N2), *mu86* mutant [CF1038, *daf-16(mu86)*] and *daf-16* OE (overexpression) [TJ356, zIs356 [*daf-16p*::*daf-16a/b*::GFP + *rol-6(su1006)*]], in the presence of Mn exposure. We found that the mutant *mu86* was more sensitive to Mn exposure; in contrast, *daf-16* OE worms were more resistant, compared with the control (Fig. 6A). Next, we crossed *mu86* mutant with MAB415 (*hpo-9* OE) worms. We found that these animals MAB456 (*hpo-9* OE, *mu86*) became more sensitive to Mn-induced lethality, with similar survival rate of *tm3719* mutants, but significantly lower than *hpo-9* OE animals (Fig. 6B), suggesting the protection of *hpo-9* OE was completely eliminated in the *mu86* mutant. Moreover, ROS levels in MAB456 animals were elevated analogous to *tm3719* animals, compared with *hpo-9* OE worms (Fig. 6C). Furthermore, in DAergic neurons, loss of *daf-16* also diminished the protection of *hpo-9* OE and resulted in more severe morphological damages, which was similar to those in *tm3719* animals (Fig. 6D). Taken together, our results indicated that DAF-16/FOXO is required for HPO-9 protection against Mn-induced toxicity in both the whole tissue and DAergic neurons. As HPO-9 is dependent on DAF-16, we assumed that it acts upstream of DAF-16.

**Figure 6.**
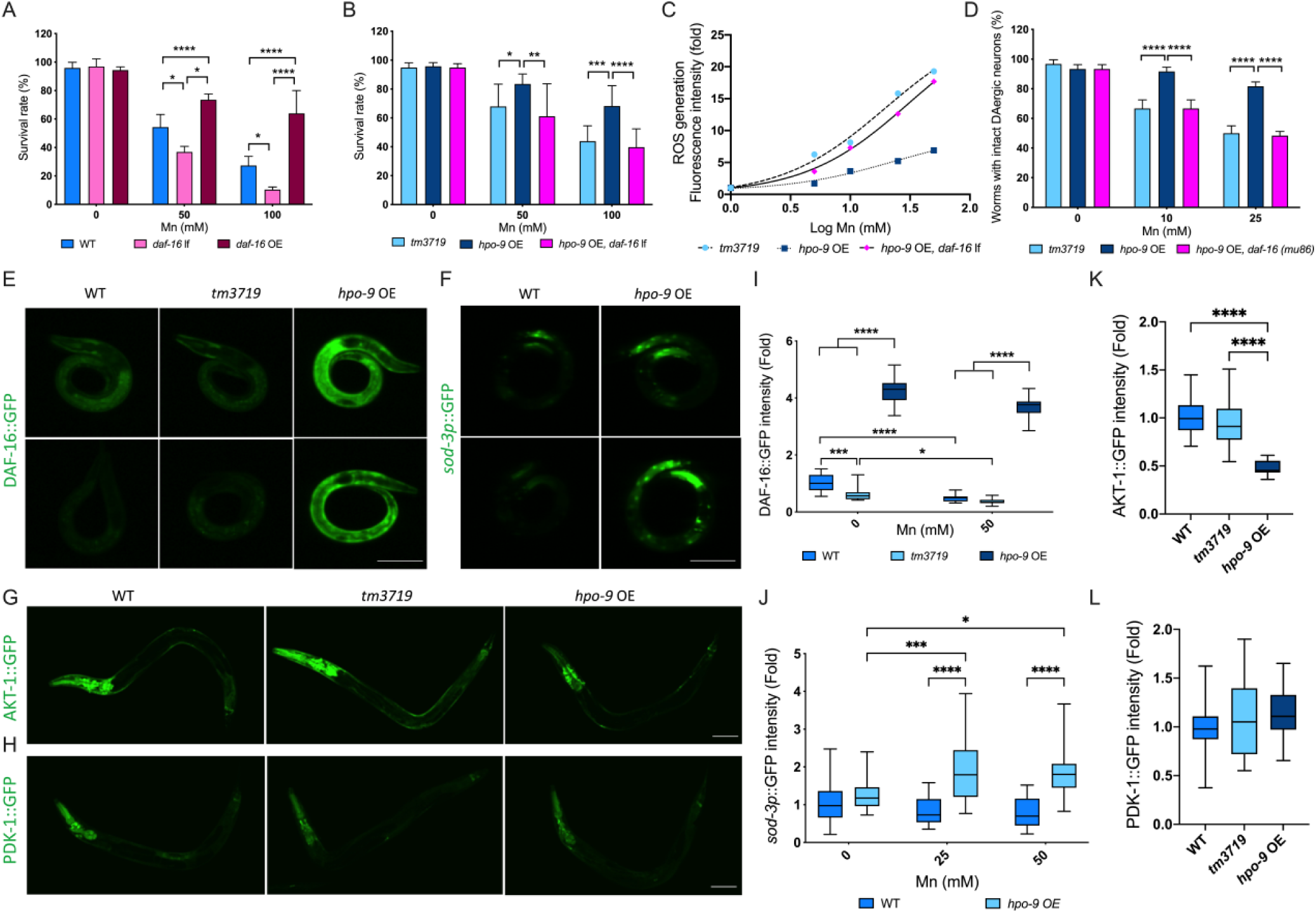
HPO-9 regulates IGF signaling. A&B Survival rates of L1s after Mn exposure. Two-way ANOVA was carried out by Graphpad Prism; *, p<0.05, **, p<0.01, ***, p<0.001, ****, p<0.0001; mean±SD (N=3). C ROS levels upon Mn exposure. Fluorescence intensities were normalized to each strain at 0 mM Mn. Two-way ANOVA by Graphpad Prism shows a significant effect between genotypes (p<0.0001); mean±SD (N=9). D Quantitative analysis of DAergic neuron morphology after Mn exposure. Two-way ANOVA was carried out by Graphpad Prism; ****, p<0.0001; mean±SD (N=3). E&F Representative images of DAF-16::GFP and *sod-3p*::GFP, respectively. Animals were synchronized at L1 stage and exposed to MnCl_2_ for 2 hours at 0 mM (upper panel) or 50 mM (lower panel), followed by 2 hours recovery before analysis. Scale bars, 50 μM. G&H Representative images of AKT-1::GFP and PDK-1::GFP in L3 animals, respectively. Scale bars, 50 μM. I&J Quantitative analysis of GFP intensity in DAF-16::GFP and *sod-3p*::GFP animals, respectively. All GFP intensities were normalized to WT animals at 0 mM Mn. Two-way ANOVA was carried out by Graphpad Prism; *, p<0.05, **, p<0.01, ***, p<0.001, ****, p<0.0001; mean±SD (n=24 animals). K&L Quantitative analysis of GFP intensity in AKT-1::GFP and PDK-1::GFP animals, respectively. All GFP intensities were normalized to WT animals. one-way ANOVA was carried out by Graphpad Prism; ****, p<0.0001; mean±SD (K, n=29 animals; L, n=36 animals).

### HPO-9 downregulates PKB/AKT level and elevates DAF-16 and downstream signaling

As the function of HPO-9 is dependent on DAF-16, it is likely that HPO-9 functions though DAF-16/FOXO signaling pathway. To study whether HPO-9 regulates DAF-16 directly, *tm3719* and *hpo-9* OE animals were crossed with TJ356 strain with a translational GFP reporter (DAF-16::GFP, Fig. 8A). We found that *tm3719* animals (MAB447) had a slight, but significant decrease in DAF-16::GFP level; surprisingly, DAF-16 level was increased to ∼4 fold in *hpo-9* OE animals (MAB444), when compared with the control (Fig. 6E&I). After Mn exposure, DAF-16 was significantly decreased in all three strains; however, the decrease in *hpo-9* OE was subtle, given it still had ∼4-fold DAF-16 levels compared with untreated control animals (Fig. 6E&I). Interestingly, there was no significant difference in DAF-16 levels between the control and *tm3719* animals after Mn exposure, suggesting that Mn decreased DAF-16 levels.

To further study whether DAF-16 is still able to activate downstream signaling upon Mn exposure, CF1553 strain (muls84[*sod-3p*::GFP + *rol-6(su1006)*]), with a transcriptional GFP reporter of *sod-3*, was crossed with *hpo-9* OE animals (MAB445). (Due to the adjacent locations of *hpo-9* gene and muls84 insertion site in the genome, we were unable to cross CF1553 with *tm3719* animals after multiple trials). No significant difference was seen in *sod-3p*::GFP levels between WT and *hpo-9* OE animals without Mn exposure; in the presence of Mn, *sod-3* level was slightly decreased in WT animals; surprisingly, *sod-3* was significantly increased in *hpo-9* OE animals (Fig. 6E&I), indicating DAF-16 was still functional and was able to respond to oxidative stress caused by Mn. In contrast, no difference was seen in *gst-4p*::GFP, a downstream target of the nuclear factor erythroid 2–related factor 2 (Nrf2/SKN-1) signaling pathway (Fig. S3).

To further identify the HPO-9 substrate in IGF signaling, proteins acting upstream of DAF-16/FOXO were investigated. Translational reporter strains expressing GFP tagged AKT-1 [GR1672 (AKT-1::GFP)] (Fig. 6G) and PDK-1 [GR1674 (PDK-1::GFP)] (Fig. 6H) were crossed with *tm3719* and *hpo-9* OE animals. Both AKT-1 and PDK-1 were highly expressed in the anterior areas of the nematode, especially the nerve ring and pharynx. We found that AKT-1 level was significantly decreased by ∼50% in *hpo-9* OE animals, but not in the *tm3719* worms (Fig. 6G&K). In contrast, PDK-1 level remained unchanged in both *tm3719* and *hpo-9* OE animals (Fig. 6H&L). AKT-1/PKB is known to phosphorylate DAF-16/FOXO and facilitate its export and degradation in the cytosol by 14-3-3 proteins (Berdichevsky, Viswanathan et al., 2006, Li, Tewari et al., 2007), consistent with our results that *hpo-9* expression increases DAF-16 levels and its downstream target SOD-3. Therefore, AKT is likely a downstream target of *hpo-9* in the IGF signaling pathway. Taken together, our results suggested that HPO-9 might specifically regulate FOXO/DAF-16 signaling, but not NRF-2/SKN-1 signaling. These data are consistent with a mechanism by which *hpo-9* downregulates AKT protein levels, in turn, increasing FOXO levels. The activation of FOXO/DAF-16 downstream target genes offers protection against Mn-induced cellular stress and neurotoxicity.

## Discussion

### Manganese as a potential environmental risk factor for RLS

The pathology of RLS is linked to Fe deficiency and dysregulation of the DAergic system. However, the etiology of RLS remains unknown. It has been reported that the level of symptoms and age of onset in RLS patients varies remarkably despite high penetrance (Dhawan, Ali et al., 2006), suggesting that environmental risk factors could be the triggering cause. Our results suggest that Mn is a potential environmental risk factor for RLS, given the strong association between BTBD9 with RLS. Loss of BTBD9/*hpo-9* in worms increased their sensitivity to Mn induced oxidative stress (Fig. 1) and DAergic neurological dysfunction (Fig. 3). In human cohort, high levels of Mn negatively regulate BTBD9 transcription in whole blood. Interestingly, an RNAseq study in human neuroblastoma cells found a significant decrease in BTBD3 (a BTBD9 paralog) after Mn exposure (p=0.03629) (Fernandes, Chandler et al., 2019). As the major risk genotypes of BTBD9 are located in the noncoding regions (Jiménez-Jiménez et al., 2018, Vilariño-Güell et al., 2009, Winkelmann et al., 2007), it is plausible that those SNPs regulate transcription and alter its mRNA levels. Our study indicates that the impact of Mn on BTBD9 gene transcription may mimic that of the risk alleles. Meanwhile, high levels of blood Mn was associated with decreased Fe levels, especially in females (Fig. 2). However, direct evidence in people with excessive Mn levels and RLS symptoms is needed to evaluate whether Mn is a causative risk factor for RLS. Due to the limited sample size in our study cohort and other RLS cohorts (Chen et al., 2020), high blood Mn levels have not been reported in those diagnosed with RLS yet.

### BTBD9 as a novel component of IGF signaling

Our results also revealed that HPO-9/BTBD9 downregulates AKT-1/PKB protein levels and upregulates DAF-16/FOXO levels, and its protection against Mn toxicity is dependent on DAF-16/FOXO signaling (Fig. 7&8). Previous *in vivo* and *in vitro* evidence support a role for Mn-dependent regulation of the IGF signaling (Bryan et al., 2020, Mooney & Green, 1989, Xu et al., 1995). In rats, Mn-deficiency has been shown to cause glucose intolerance and reduced insulin production (Baly, Schneiderman et al., 1990). whilst excessive Mn increased *IGF1* mRNA and IGFR protein levels (Hiney, Srivastava et al., 2011). In *C. elegans*, Mn exposure has been shown to decrease FOXO and its downstream target SOD-3 protein levels, while increasing AKT levels (Ávila, Somlyai et al., 2012). In addition, AKT and SGK (serum and glucocorticoid-regulated kinase) loss-of-function mutants were shown to be more resistant to Mn-induced toxicity (Peres et al., 2018). Collectively, these *in vivo* studies support a functional link between Mn biology and IGF signaling. In mouse neuronal and human stem cell-based models, Mn-dependent insulin/IGF signaling has been shown to be dependent on bioavailable Mn acting directly at the level of the IR/IGFR (Bryan et al., 2020). Further, *in vitro* biochemical evidence indicates that the insulin receptor (IR) and IGFR exhibit increased kinase activity in the presence of Mn versus magnesium as the kinase co-factor (Mooney & Green, 1989, Xu et al., 1995). Our present studies provide evidence of additional putative mechanisms by which Mn may regulate IGF signaling: Mn exposure a) downregulates *hpo-9*/BTBD9 levels (Fig. 2); or b) decreases *hpo-9*/BTBD9, resulting in lower DAF-16/FOXO levels (Fig. 8); c) which inhibits activation of downstream target genes.

It has been reported that BTBD9 functions as an adaptors for the cullin-3 (Cul-3) class of E3 ubiquitin ligases (Freeman et al., 2012, Li, Zhang et al., 2020). Our results showed that expression of HPO-9 significantly decreases PKB/AKT-1 level (Fig. 8M-O&S) and dramatically increases FOXO level (Fig. 8A-F&Q). Thus, it is reasonable to assume that BTBD9 may promote protein degradation and/or dephosphorylation of AKT, which eventually upregulates FOXO protein levels. We assume BTBD9 may have similar function as phosphatase and tensin homolog (PTEN) and the serine/threonine phosphatase 2 (PP2A) to counteract IGF signaling. In addition to FOXO, AKT also directly or indirectly regulates several other transcription factors [transcription factor EB (TFEB) (Palmieri, Pal et al., 2017), tumor protein p53 (Abraham & O’Neill, 2014) and nuclear factor kappa-light-chain-enhancer of activated B cells (NF-κB) (Bai, Ueno et al., 2009)], as well as two multifunctional downstream signaling nodes-glycogen synthase kinase 3 (GSK3) and mammalian target of rapamycin complex 1 (mTORC1) (Manning & Toker, 2017). Through these targets, AKT is involved in multiple signal transduction pathways and regulates expression of genes in stress response, autophagy and lysosome, cell survival and apoptosis, inflammation and immune response, etc. As Mn exposure activates AKT and downregulates *hpo-9*/BTBD9 mRNA level (Fig. 2), it is possible that *hpo-9*/BTBD9 is negatively regulated by AKT. FOXO has important neuroprotective function as it regulates neuronal autophagy and oxidative stress (McLaughlin & Broihier, 2018). Therefore, it is noteworthy that FOXO interacts with TFEB and regulates mitochondrial uncoupling proteins (UCPs) (Liu, Tao et al., 2016). Previously we have shown that Mn exposure downregulates nuclear localization of TFEB and suppresses autophagic-lysosomal degradation of unhealthy mitochondria (Zhang, Yan et al., 2020). Here we showed loss of *hpo-9* resulted in significant accumulation of abnormal mitochondria and it was worsened by Mn exposure (Fig. 1I-L), which revealed the important role of BTBD9 in energy production, especially in the nervous system. Given that *hpo-9* OE increases FOXO and *sod-3* levels, and protects against Mn-induced DAergic neurodegeneration, DAergic neurons lacking BTBD9 are less capable to produce ATP and defend oxidative stress, and thus more susceptible to intrinsic/extrinsic DAergic insults, which might contribute to the etiology of RLS associated BTBD9 risk alleles. Together, our study sheds new light on the links between Mn, BTBD9, FOXO and mitochondrial homeostasis. Last, but not least, as BTBD9 is able to regulate AKT/PKB, we expect that this protein has a boarder function in addition to mitigating against oxidative stress caused by various environmental toxins. However, further studies are needed to investigate the specific interaction between BTBD9 and AKT/AKT-regulated signaling pathways and confirm the role of BTBD9 in more general oxidative stress.

## Materials and Methods

### Plasmid constructs

*hpo-9* coding sequence (∼2.1kb, with C-terminal FLAG tag) and *hpo-9* promoter sequence (∼1.5kb upstream of the start codon) was PCR amplified using from genomic DNA isolated from wildtype (WT) N2 worms. The human BTBD9 cDNAs (with C-terminal FLAG tag) were amplified from cDNAs provided by M. Diana Neely (Vanderbilt University, Nashville, TN). The GFP tag was fused to *hpo-9* and BTBD9 C-terminal by fusion PCR. The transcriptional GFP reporter *hpo-9p::*GFP*::unc-54* 3’UTR was created by fusing *hpo-9* promoter sequence, GFP coding sequence and *unc-54* 3’UTR together. Using Gateway recombinational cloning (Invitrogen), the above PCR products were recombined with the pDONR221 vector to create pENTRY clones. For somatic expression in worms, *hpo-9* and BTBD9 pENTRY constructs were then recombined into pDEST-*eft-3* vector, under the promoter from eukaryotic translation elongation factor (*eft-3*) gene. For expression in DAergic neurons, the pENTRY constructs were recombined into pDEST-*dat-1* vector, under the promoter from the DA transporter (*dat-1*) gene. For body wall muscle expression, HPO-9::GFP pENTRY constructs were then recombined into pDEST-*unc-54* vector, under the promoter from a muscle myosin class II heavy chain (*unc-54*) gene. These plasmids were then used to create transgenic worms. For expression in Nuro2A cells, *hpo-9* and BTBD9 (without stop codon) pENTRY constructs were recombined into pcDNA-DEST47 vector for later transformation.

### C. elegans strains

Nematodes were grown and maintained using standard procedures.(Brenner, 1974) The strain MAB200 carrying a *hpo-9* mutant (allele *tm3719* with 761bp deletion) was ordered from National BioResource Project: *C. elegans* in Japan. Strains CF1038, CF1553, CL2166, GR1672, GR1674, OH7193, SD1347 and TJ356 were ordered from *C. elegans* Genetic Center (CGC). All strains and genotypes are listed in Table S1. Transgenic worms were created using microinjection as previously described (Mello, Kramer et al., 1991). Plasmids were injected at a concentration of 50 mg/ml. For whole worm expression, *eft-3*::*hpo-9* or BTBD9 alone were co-injected into tm3719 strain with pG2M36 (*myo-3*::dsRed) and pBCN27-R4R3 (*rpl-28*::PuroR, Addgene), which were used as the selective markers for transformation. For expression in DAergic neurons, *dat-1p*::*hpo-9* or BTBD9 alone were co-injected into MAB300 [*dat-1*::GFP*(vtIs1) V*, *smf-2*(*gk133*) *X*] strain with *elt-2*::mCherry and pBCN27-R4R3. The *hpo-9* transcriptional GFP reporter was generated by injecting *hpo-9p::*GFP::*unc-54* 3’UTR, pRF4 and pBCN27-R4R3 into OH7193 (otIs181 [*dat-1p*::mCherry; *ttx-3p*::mCherry], *him-8(e1489)* IV) worms. For the translational reporter, *unc-54p*::HPO-9::GFP was co-injected with pG2M36 (*myo-3*::dsRed) into MAB200 worms. For each injection mixture, at least three stable lines were generated and analyzed. Representative lines were selectively integrated by using UV irradiation using a Spectroline UV crosslinker with an energy setting of 500 mJ/cm2.

### Metal exposure

Worms were synchronized at Larva 1 (L1) or Larva 4 (L4) stages as per experimental needs. MnCl_2_, FeCl_2_, CuCl_2_ and ZnCl_2_ exposures were performed as previously described (Chen, Bowman et al., 2015, Leyva-Illades et al., 2014). As the *C. elegans* cuticle is a barrier to permeation, Mn doses used in the worm are generally higher than those in mammals (Nass et al., 2002, Rand & Johnson, 1995), but the intracellular Mn levels are similar (Benedetto, Au et al., 2010).

### Lethality analysis

Survival assay was performed as previously described (Leyva-Illades et al., 2014). Experiments were performed in at least three independent replicates.

### RNA interference (RNAi) in *C.elegans*

RNAi by feeding was performed as previously described (Chen, Burdette et al., 2010), except that a RNAi sensitive strain GR1373 (*<u>eri-1</u>(<u>mg366</u>*)) was used in this study. Worms were first synchronized at L1 stage and then grown on RNAi plates with RNAi bacteria for 2 day until they reached young adult stage. The adults were then exposed to MnCl_2_ for 2 hours, and the survival rate were analyzed one day later. The bacterial clone producing dsRNA for *hpo-9* was obtained from the Ahringer RNAi library (Source BioScience). The control bacteria contained the empty RNAi expression vector pL4440.

### Measurement of intracellular reactive oxygen species (ROS)

After Mn exposure, L1 nematodes were washed and resuspended in M9 buffer. ∼500 L1s (in 50μl M9) were then transferred to 96-well plates, with 50μl of 50μM fluorescent probe 2′,7′-dichlorofluorescein diacetate (DCFDA) (at a final concentration of 25 μM).(Yoon et al., 2018) Worms were gently shaken for 2 hours, then the fluorescence intensity was quantified by a plate reader (FLUOstar OPTIMA, BMG LABTECH) at an excitation wavelength of 490 nm and an emission wavelength of 510–570 nm.

### Mitochondrial morphology

Nematodes with mitochondria-localized GFP in body wall muscle were used to assess alterations in mitochondrial morphology. Synchronized L4 larva were treated with MnCl_2_ for 2 hours, followed by 2-hr recovery. Worms were then mounted onto a 4% agarose pad. Images were taken immediately on a Leica SP8 Confocal Microscope with excitation/emission wavelengths at 488/520 nm. Mitochondrial morphology was assessed in at least twenty animals for each condition, and blindly scored. The morphological categories of mitochondria were defined according to Momma, Homma et al. (2017) with slight modification: (1) tubular: a majority of mitochondria were interconnected and elongated like tube shape; (2) intermediate: a combination of interconnected and fragmented mitochondria; (3) fragmented: a majority of round or short mitochondria in the image taken.

### DAergic neurodegeneration

The Morphology of DAergic neurons was examined as previously described (Leyva-Illades et al., 2014). For each strain and condition, at least 20 animals were quantitated in three independent replicates.

### Confocal microscopy

Fluorescent images were taken by confocal microscopy as previously described (Benedetto et al., 2010). Worms expressing DAF-16::GFP and *sod-3p::*GFP were analyzed at L1 stage; worms expressing AKT-1::GFP and PDK-1::GFP were analyzed at L3 stage.

### Formaldehyde induced fluorescence (FIF)

After Mn exposure, L1 animals were recovered on seeded NGM plates for 2 hours. Next, FIF was performed as previously described (Lints & Emmons, 1999, Rivard et al., 2010). Fluorescent intensity was quantified by ImageJ. For each experiments, ten individual DAergic neurons (CEPs) were measured in three independent replicates.

### Basal slowing response

The behavioral assay was performed as previously described (Sawin et al., 2000), except that the worms were exposed to Mn at L1 stage and assayed at L4/young adult stage.

### Quantitative real time PCR (qPCR)

qPCR was applied to determine the expression levels of target genes using TaqMan Gene Expression Assays (Applied Biosystems). The gene *ama-1* (Ce02462726_m1), encoding an RNA polymerase II large subunit was used as the control. The *hpo-9* (Ce02477190_g1) probe with Exon 4-5 boundary was able to detect the mRNAs from both the wild type N2 and the mutant *tm3719* (deletion in Exon 2) animals.

### Semi-quantitative RT-PCR

Semi-quantitative RT-PCR was performed on worms to validate the expression of *hpo-9(tm3719)* allele and transgenes. RNA isolation was performed as described (Chen et al., 2010). To detect the truncated *tm3719* allele, primers 5’-ATGAGCGATAACCATGCTTTTGG-3’ and 5’-TTATTTTATGGCAATTGGAACGTTTG-3’ are used. To detect FLAG tagged transgenes (*hpo-9* and *BTBD9*) in DAergic neurons, the following primer pairs were used: for *hpo-9*::FLAG, 5’-ATGAGCGATAACCATGCTTTTGG -3’ and 5’-TCACTTGTCATCGTCGTCCTTGTAGTC-3’; for *BTBD9*::FLAG, 5’-ATGAGTAACAGCCACCCTCTTC -3’ and 5’-TCACTTGTCATCGTCGTCCTTGTAGTC-3’. For the *ama-1* (RNA polymerase II large subunit) loading control, the forward primer was 5’-GCTACTCTGGCAAGACGTG-3’, and the reverse primer was 5’-CGAGCGCATCGATGACCC-3’.

### Human blood studies

Individuals were randomly selected from the follow-up of Mn-exposed workers healthy cohort (MEWHC) in 2020, as previously described (Ge, Liu et al., 2020, Zhou, Ge et al., 2018). Whole blood samples were collected in tubes containing ethylenediamine tetraacetic acid (EDTA) anticoagulant from the participants after overnight fasting, and stored at −80°C until RNA extraction and metal analysis. The present study was approved by the Medical Ethics Committee of Guangxi Medical University. Concentrations of Mn and Fe were quantified by inductively coupled plasma-mass spectrometry (ICP-MS, Thermo Scientific) as previous described (Hou, Huang et al., 2019), except for the following details. Briefly, diluent containing 0.1% nitric acid (Fisher Scientific), 0.5% 1-Butanol (Acros) and 0.1% Triton ™ X-100 (Sigma) was used to dilute whole blood samples. Total RNA was extracted from 250 μL of whole blood using TRIzol LS Reagent (Invitrogen). A total of 300 ng of RNA was used to obtain complementary DNA (cDNA) by reverse transcription. Real-time quantitative PCR (RT-qPCR) used 2×RealStar Green Fast Mixture (GenStar) with a Real-time PCR Detection System (LightCycler96, Roche). The following primers were used in the amplification: BTBD9: forward 5’-AGCCTGCCTCCTTCATCCGTATC-3’ and reverse 5’-CTCTGCTGCTCTGGACACTCAAAG-3’; Actin: forward 5’-CATGTACGTTGCTATCCAGGC-3’ and reverse 5’-CTCCTTAATGTCACGCACGAT-3’.

### Neuro2A cell culture

Neuro2a cells were maintained in an incubator with the supplementation of 5% CO2 at 37 °C. The normal culture medium is Dulbecco’s modified Eagle medium (DMEM) containing 10% (v/v) fetal bovine serum (FBS, Gibco), 10 U/ml penicillin and 100 μg/ml streptomycin. For transformation, cells were placed in a 6-well plate and incubated in DMEM with 10% FBS until they reached to ≥80% of confluency. Cells were then transfected with pGFP, pGFP-HPO-9 or pGFP-BTBD9a plasmid using TransIT®-LT1 (Mirus). The cells were incubated with transfection complex for 24 hours at incubator with the supplementation of 5% CO2 at 37 °C. After transfection, the HPO-9::GFP and BTBD9a::GFP cells were placed in 96-well plates and treated with 0, 0.125, 0.25 and 0.5 mM/L MnCl2 for 24 hours, respectively. Cellular viability was measured using MTT assay. The absorbance was measured at 490 nm with a plate reader (FLUOstar OPTIMA, BMG LABTECH).

### Statistical analysis

Statistical analyses were conducted in GraphPad Prism version 8 (GraphPad Software Inc.), and results were expressed as mean ± standard deviation (SD). Student’s *t-*test or one-way analysis of variance (ANOVA) were performed to analyze data with one factor. Data with two factors were analyzed by two-way ANOVA followed by Sidak’s multiple comparisons test when main effects were observed. Differences were considered statistically significant if p-value was <0.05.

## Acknowledgements

We thank the patients and families for their participation in this study. We thank the *Caenorhabditis* Genetics Center (CGC) and Shohei Mitani for providing the *C. elegans* strains. We also thank Dr. M. Diana Neely for the human cDNAs. This study was supported by National Institute of Environmental Health Sciences grants NIEHS R21ES031315, NIEHS R01ES10563 and NIEHS R01ES07331. We thank further the German Research Foundation (DFG) for the financial support of the DFG Research Unit TraceAge (FOR 2558), National Natural Science Foundation of China [grant number 81860573 and 82073504] and Guangxi Natural Science Foundation for Innovation Research Team [grant number 2017GXNSFGA198003].

## Author Contributions

P.C., F.Z., S.L., H.C., X.Y. Y.L., B.Y., K.L., T.K., J.B., T.S. were involved in execution of experiments and analysis of data. P.C., A.B.B. and M.A. were responsible for conception, organization and drafting of the manuscript.

## Expanded Figure Legends and Tables

**Figure S1.**
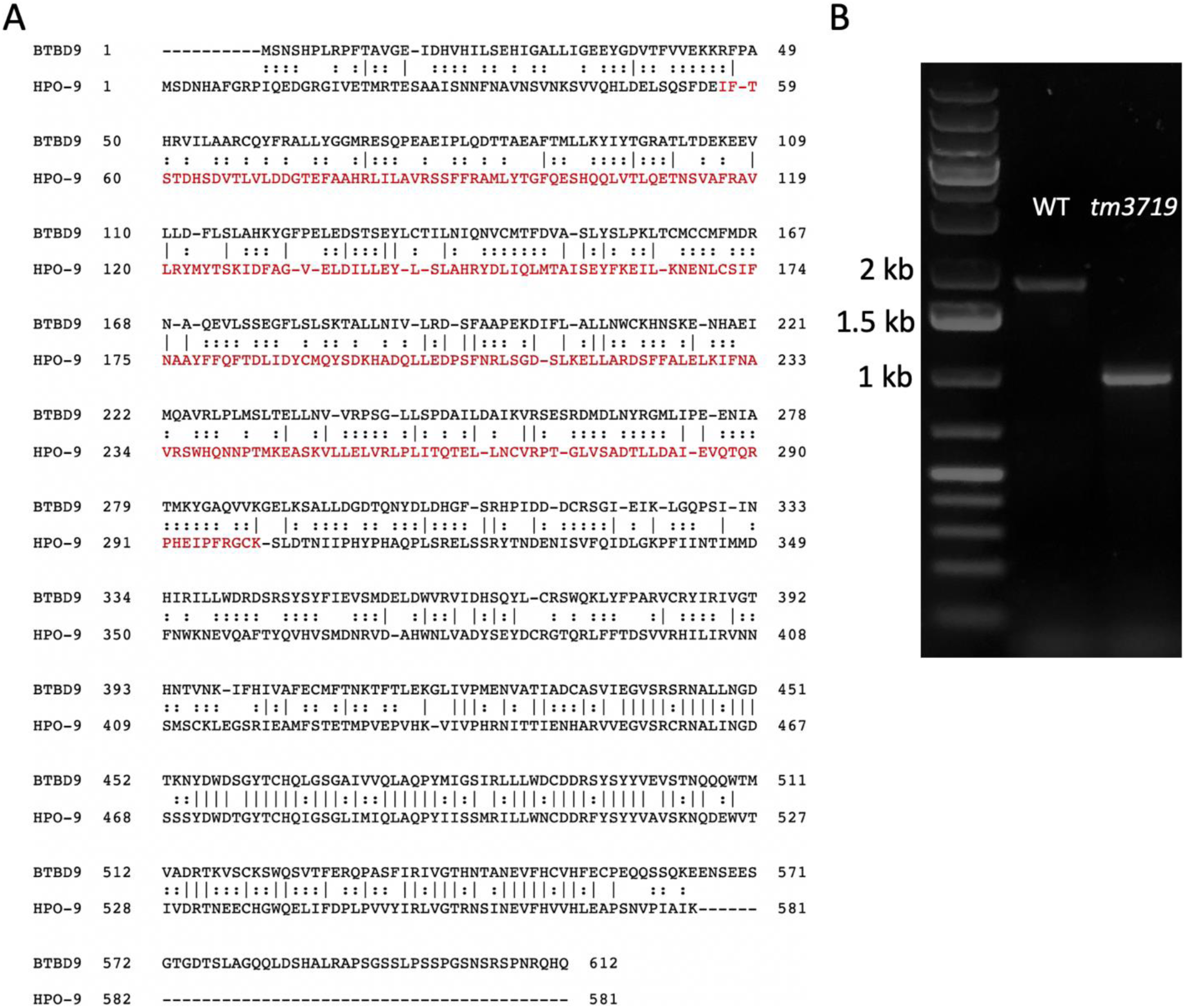
The *C. elegans* BTBD9 homolog HPO-9 and its mutant allele *tm3719*. A Protein alignment of BTBD9 isoform a and HPO-9. The sequence similarity is ∼75%. Red, amino acids deletion in *hpo-9(tm3719)* allele. B *hpo-9* gene products in the control (N2 WT) and *tm3719* mutant worms.

**Figure S2.**
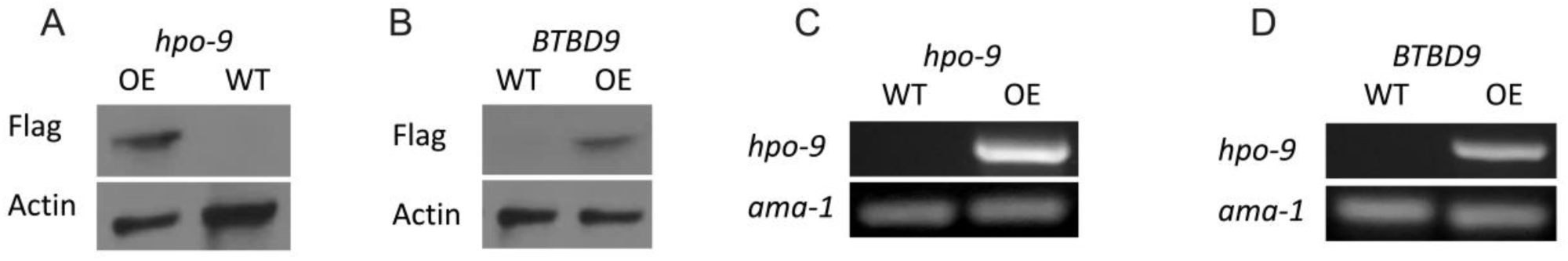
Expression of transgenes. A&B Detection of HPO-9::FLAG and BTBD9a::FLAG by Western blot in MAB415 and MAB425 animals. C&D Detection of HPO-9::FLAG and BTBD9::FLAG in MAB414 and MAB405 animals by semi-quantitative RT-PCR. OE, overexpression strains; WT, N2 strains.

**Figure S3.**
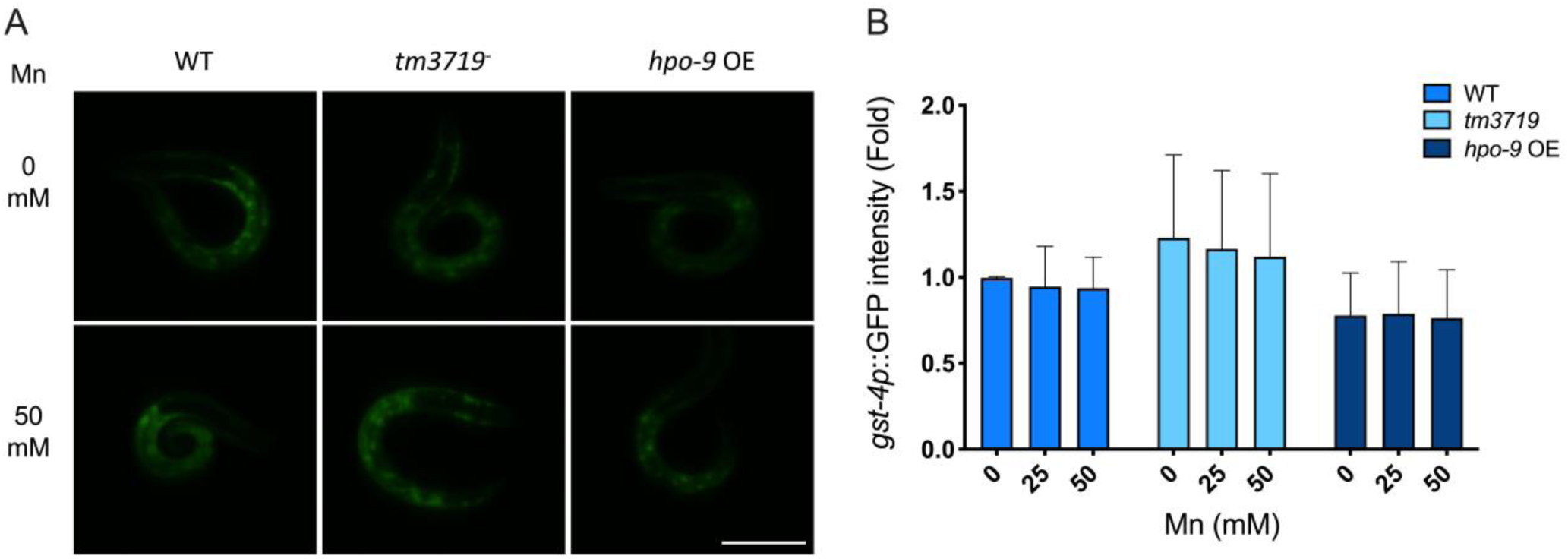
HPO-9 does not regulate *gst-4p*::GFP. A Animals were synchronized at L1 stage and exposed to MnCl_2_ (0, 25 and 50 mM) for 2 hour. Images were taken by confocal microscopy and GFP intensities were analyzed using ImageJ; GFP intensity was normalized to the WT animals at 0 mM Mn. Scale bar, 50 μM. B Quantitative analysis of GFP intensity. For each experiment, ∼15 animals were analyzed. Two-way ANOVA as carried out by Graphpad Prism; mean±SD (N=3).

**Table S1.**
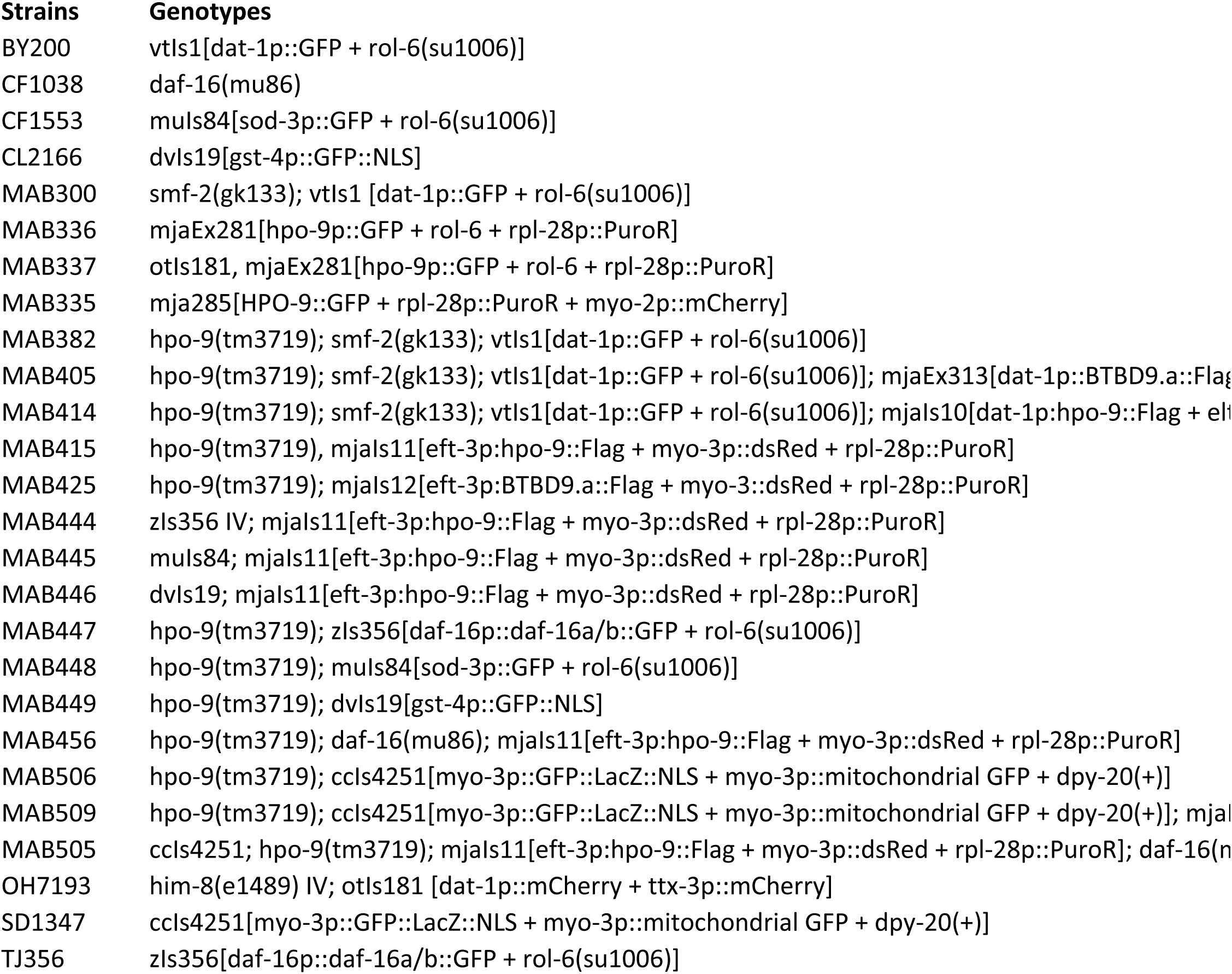
List of strains and genotypes.

